# When can fitness epistasis be ignored in a polygenic trait at equilibrium?

**DOI:** 10.1101/2023.01.25.525607

**Authors:** Archana Devi, Kavita Jain

**Author notes:** (A. Devi). (K. Jain). https://ufind.univie.ac.at/en/person.html?id=1010614 (A. Devi); https://www.jncasr.ac.in/faculty/jain (K. Jain).

## Abstract

Although many phenotypic traits are determined by a large number of genetic variants, the behavior of allele frequencies in a polygenic trait is not completely understood. The problem is especially challenging when the quantitative trait of interest is under epistatic selection as the allele frequency at a locus is affected by those at other loci. Here, we consider a panmictic, diploid finite population evolving under stabilizing selection and symmetric mutations when the population is in linkage equilibrium. In the stationary state, using a diffusion theory, we calculate the marginal distribution of allele frequency, and find parameter regimes where fitness epistasis can not be ignored for an accurate description of the frequency distribution. For such parameters, the mean deviation in the phenotypic optimum and genic variance are, however, found to be well captured even when epistatic interactions are neglected. Thus, while the presence of epistasis may not be evident in phenotypic quantities, it can strongly affect the allele frequency distribution. We also find that the allele frequency distribution at a locus is unimodal if its effect size is below a threshold effect and bimodal otherwise; these results are the stochastic analog of the deterministic ones where the stable allele frequency becomes bistable when the effect size exceeds a threshold. Our analytical results are verified against Monte Carlo simulations and numerical integration of a Langevin equation.

## 1. Introduction

The evolutionary dynamics of complex phenotypic traits have traditionally been studied in the framework of quantitative genetics that deals with the summary statistics of a trait; however, a detailed understanding of the genetic basis of phenotypic variation is crucial for predicting the evolutionary outcomes of quantitative traits such as complex diseases (García-González and O’Reilly, 2026). Population genetics focuses on the genetic details of the phenotypic trait as the allele frequencies underlying a trait evolve under the joint action of evolutionary forces such as mutation, selection, random genetic drift and gene flow, and informs us on the evolution and maintenance of phenotypic variation in a population and between populations and species. Due to the advancement of genome-wide association studies (GWAS) in recent years, much information about the genetic architecture of phenotypic traits has increasingly become available, and it is now known that the phenotypic variation depends on the number of genetic variants underlying a phenotype, the effect size and frequency of these variants, the interaction between these genetic variants and with the environment, etc. (Visscher, Wray, Zhang, Sklar, McCarthy, Brown and Yang, 2017; Timpson, Greenwood, Soranzo, Lawson and Richards, 2018). For these reasons, much work in the last two decades has focused on understanding quantitative traits at a genetic level and connecting these insights to the phenotype (Rockman, 2012; Jain and Stephan, 2017a; Sella and Barton, 2019; Barghi, Hermisson and Schlötterer, 2020).

The evolution of a quantitative trait in terms of the underlying allele frequencies is also theoretically challenging (Bürger, 2000; Walsh and Lynch, 2018) since stochastic models of polygenic adaptation are described by high-dimensional Fokker-Planck equations, and the phenotypic selection is typically epistatic, that is, it is nonlinear in the trait value and therefore can not be expressed as the sum of fitness of individual phenotypic values. Several recent studies have shown that phenotypic traits such as birth weight in humans are under stabilizing selection which has a tendency to reduce phenotypic variation in a population as the fitness of a trait value decreases with increasing deviation from the phenotypic optimum (Robertson, 1956; Kingsolver, Hoekstra, Hoekstra, Berrigan, Vignieri, Hill, Hoang, Gibert and Beerli, 2001; Sanjaka, Sidorenkoc, Robinson, Thornton and Visscher, 2018; de Villemereuil, Charmantier, Arlt, Bize, Brekke, Brouwer, Cockburn, Côté, Dobson, Evans, Festa-Bianchet, Gamelon, Hamel, Hegelbach, Jerstad, Kempenaers, Kruuk, Kumpula, Kvalnes, McAdam, McFarlane, Morrissey, Pärt, Pemberton, Qvarnström, Røstad, Schroeder, Senar, Sheldon, van de Pol, Visser, Wheelwright, Tufto and Chevin, 2020; Sodeland, Jentoft, Jorde, Mattingsdal, Albretsen, Kleiven, Wårøy Synnes, Espeland, Olsen, Andrè, Stenseth and Knutsen, 2022). If the population is initially far from the optimum, one can describe the evolutionary dynamics assuming directional selection which, however, lacks epistasis (Götsch and Bürger, 2024). But at large times, when the population is close to the optimum or has reached mutation-selection-drift equilibrium, the epistatic nature of stabilizing selection - both at phenotypic level as also at genetic level due to phenotype-genotype map - comes into play, and it is important to understand if and how these epistatic interactions affect the allele frequency distribution and the phenotypic quantities.

Here, we consider a polygenic trait evolving under stabilizing selection and symmetric mutations in a panmictic finite population under linkage equilibrium, and focus on its equilibrium properties. Due to stabilizing selection, the allele frequencies interact with each other and evolve such that the trait value approaches the optimum. Although the joint distribution of allele frequencies in the stationary state is known (Wright, 1937; Kimura, 1964), the distribution is not transparent enough to give an insight into the genetic architecture of the phenotypic trait. In this study, we obtain the marginal distribution of allele frequencies and study its properties in detail. Our main conclusion is that for a polygenic trait controlled by a large number of loci, as is relevant in many biologically realistic scenarios, the allele frequency distribution can be accurately described ignoring epistatic interactions, provided selection is sufficiently strong. But for weak to moderate selection, this is possible under certain conditions on mutation and selection parameters and if the locus under consideration has sufficiently small effect size. However, irrespective of selection strength, the genic variance is found to match well with Bulmer’s expression (Bulmer, 1972) that has been obtained on neglecting epistasis. Our work also extends the previous theory for infinitely large populations (de Vladar and Barton, 2014; Jain and Stephan, 2017b) to large but finite populations.

## 2. Model

We consider the classic Latter-Bulmer model (Latter, 1960; Bulmer, 1972) that describes a panmictic, diploid population of size *N* characterized by a single polygenic trait which is controlled by *L* diallelic loci. The phenotype-genotype map is assumed to be additive (Hill, Goddard and Visscher, 2008), and the trait value 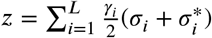 where *σ*_*i*_, 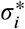, respectively, denote the maternal and paternal contribution to the trait of an individual and take the value 1(−1) if mutant (wildtype) allele is present at the *i*th locus. Here the effect size of either allele is *γ*_*i*_, and is chosen independently of the effect size at other loci from a common distribution *p*(*γ*) which is assumed to be an exponential distribution with mean 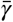 (Mackay, 2004; Goddard and Hayes, 2009).

The phenotypic trait evolves under stabilizing selection (Robertson, 1956; Kingsolver et al., 2001; Sanjaka et al., 2018; de Villemereuil et al., 2020; Sodeland et al., 2022) with the phenotypic fitness function,

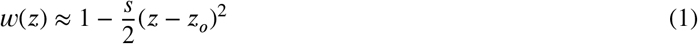

which decays quadratically away from a (time-independent) optimal trait value *z*_*o*_, and where *s* denotes the strength of selection. We also assume that the mutation between wildtype and mutant allele occurs at an equal rate *μ* at each locus.

The dynamics follow a classical Wright-Fisher model where genotypes of the individuals undergo random genetic drift, selection, recombination, mutations, and random matings with other individuals to form the next generation - while selection, recombination, and mutation are deterministic forces, the allele frequencies fluctuate in a finite population due to random genetic drift. At large times, the population always reaches a steady state under the joint action of these biological forces, and here we focus on the resulting equilibrium state. Under linkage equilibrium (see Appendix A1), the genotype frequencies can be expressed as the product of allele frequency at each locus in a fully recombining population; however, the allele frequencies do not evolve independently due to the indirect epistatic interaction imposed by the fitness function (1).

In the following sections, the model described above is studied analytically using a diffusion theory, and numerically using Monte Carlo (MC) simulations and by integrating a Langevin equation using Euler-Maruyama (EM) method. As discussed in Sec. 3 and Sec. 4.4, for a population in linkage equilibrium, the MC simulations are accurate for low mutation rates, but EM method works better for high mutation rates. For this reason, below we describe both methods.

## 3. Methods

The model dynamics implemented in MC simulations are depicted in Fig. 1. We consider a population of *N* individuals, each having two chromosomes with *L* sites and assign an effect value *γ* to each locus (the effect sizes are exponentially distributed unless stated otherwise). Then each locus of a chromosome is multiplied by +1 or −1 with its effect size, where +1 or −1 are chosen with equal probability using a uniform random number generator. The trait value of each individual is half of the sum of its two chromosomal values. The population evolves in discrete and nonoverlapping generations through the following steps that are performed at each generation:

(i) Random genetic drift: We choose individuals randomly from the population using a uniform random number generator. However, the viability of a particular individual depends on its relative phenotypic fitness keeping the total number of individuals constant every generation.
(ii) Selection: Each individual is assigned a fitness according to (1). We weight the individual fitness with the maximum fitness in the population since the viability of the individual depends on its relative fitness in the population rather than its absolute fitness. We select the individual if a uniform random number is less than its relative fitness.
(iii) Recombination: Two chromosomes of the selected individual recombines according to the recombination parameter. Since we assume free recombination in this study, we find the recombination break points by drawing a Bernoulli-distributed random number with probability one half at every locus. After finding all the breakpoints, we exchange the genetic material of the two chromosomes at each recombination breakpoint. After the recombination event, we keep either of the recombined chromosomes with equal probability and discard the other one for every individual. This process continues until we select 2*N* chromosomes.
(iv) Mutation: We change the sign of the effect size of each locus if a standard uniform random number is less than the mutation probability.
(v) Random mating: We get *N* offspring for the next generation by pairing chromosomes randomly from the 2*N* chromosomes.

**Figure 1.**
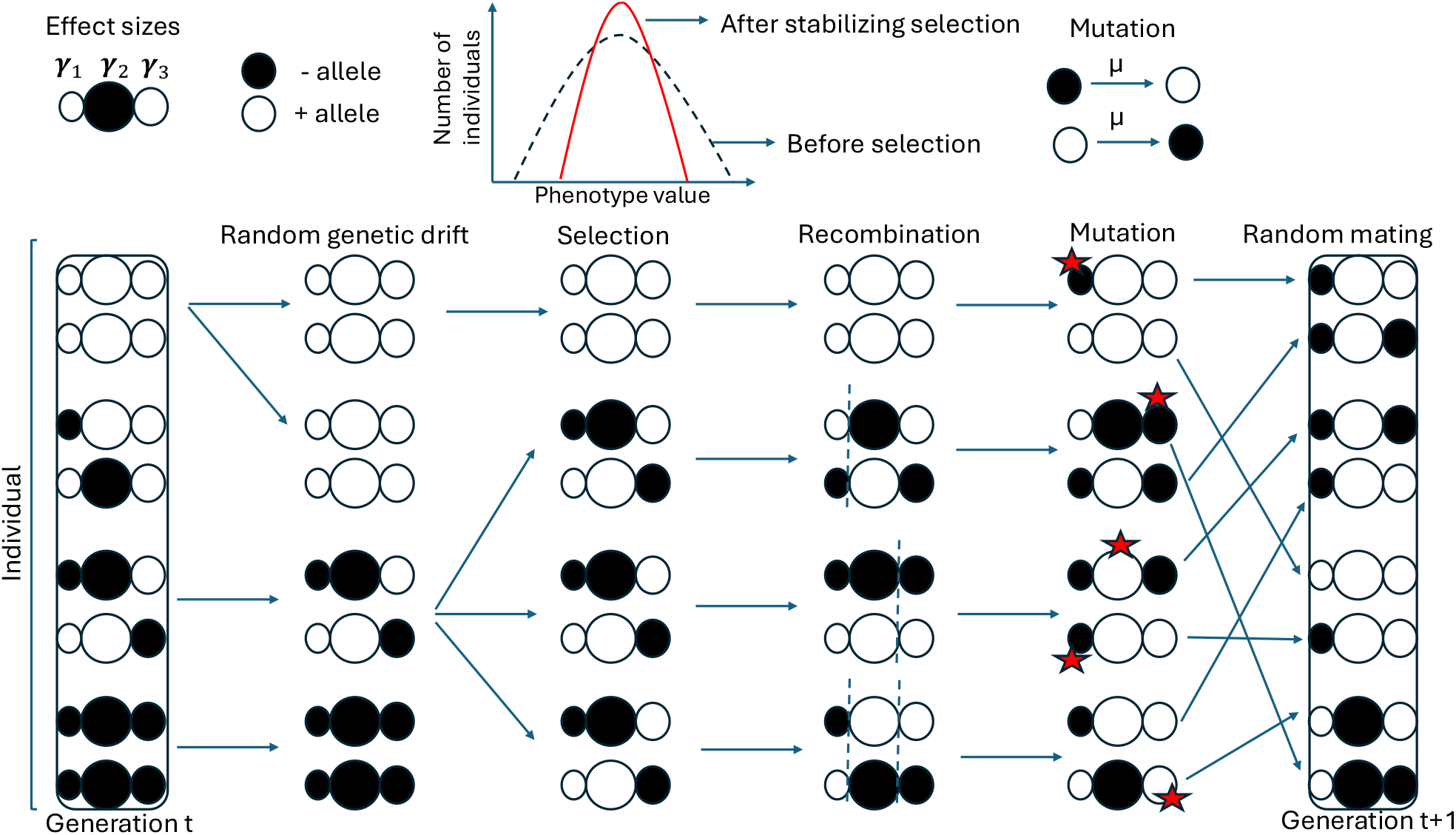
Diagram depicting how each evolutionary force shapes the genotypic configuration of individuals in a Wright-Fisher process implemented in Monte Carlo simulations. Here, we consider *N* = 4 diploid individuals with *L* = 3 loci to show the simulation steps from generation *t* to *t* + 1.

As shown in Appendix A2, under linkage equilibrium, the above model can be described using a diffusion theory. Following Itô prescription, corresponding to the Fokker-Planck equation (A2.1), the Langevin equation for the allele frequency *x*_*i*_ at the *i*th locus is given by [refer to Sec. 4.3.5, Gardiner (1997)]

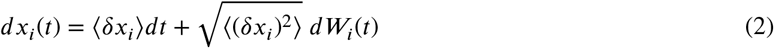

where, as discussed in Appendix A2,

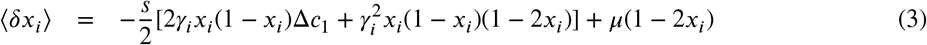

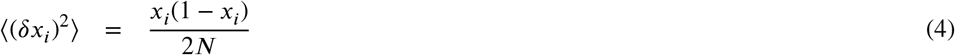

and *dW*_*i*_ is the Wiener process. The Langevin equation (2) is integrated numerically using the EM method (Higham, 2001). Dividing the time *t* into *t*/*δt* intervals of equal length *δt*, the allele frequency at the *i*th locus is updated as (Sec. 4.3.1, Gardiner (1997))

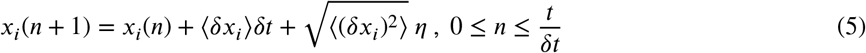

where *η* is a random variable chosen independently at each time step from a normal distribution with mean zero and variance *δt*. In all the numerical data presented using the above equation, the population starts with allele frequency one half at every locus, and time step size *δt* = 0.1.

For either method, after a burn-in period of 10*N* generations, the data in the stationary state are obtained by averaging over 10^5^ −10^8^ generations, depending on the parameters. Although the EM method is computationally faster and easier, as shown in Fig. 6, it fails to accurately capture the genic variance (see Appendix A1 and Sec. 4.4) for low mutation rates. The EM method is designed to simulate continuously fluctuating, large-population, and can not correctly model the small population dynamics or low mutation rate where fixation or loss of alleles occur (Hutzenthaler, Jentzen and Kloeden, 2011). On the other hand, the MC simulations which are computationally expensive for large populations with large number of loci, are accurate for low mutation rate; however, the linkage equilibrium gets distorted when both mutation and selection are strong in a finite population which generates (and maintains) non-random associations between loci faster than the maximum recombination probability equal to one half can dissipate them. These non-random associations prevent the population from reaching the perfect linkage equilibrium in MC simulations, but the EM method uses the Langevin equation which is derived assuming linkage equilibrium. For this reason, we use the appropriate method to obtain the numerical results discussed below.

## 4. Steady state distribution

In the steady state and under linkage equilibrium, the exact joint distribution of mutant allele frequencies has been obtained in the framework of diffusion theory, and is given by (Kimura, 1964) [also see Appendix A2]

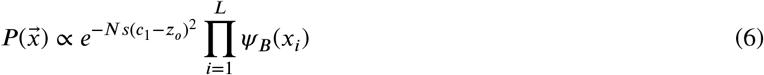

where,

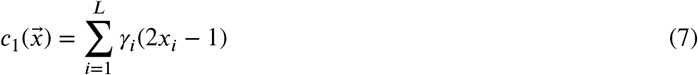

is the population-averaged phenotype, and

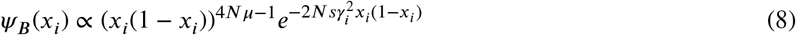

Note that due to linkage equilibrium, although the population dynamics at each locus occur independently of the other loci, the distribution (6) is not a product measure; this is because due to epistasis, the selection (1) is nonlocal and depends on all loci through the global variable 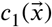.

### 4.1. Marginal distribution for a large number of loci

To understand the allele frequency distribution at a locus, we now consider the marginal distribution *ψ* (*x*_*i*_ ) of the frequency at the *i*th locus which can be obtained by integrating the joint distribution 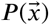 over all the allele frequencies except *x*_*i*_ and using the central limit theorem. For a large number of loci, as shown in Appendix A3, the single-locus distribution can be approximated by

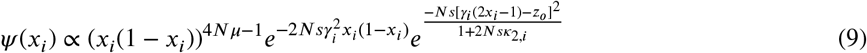

where,

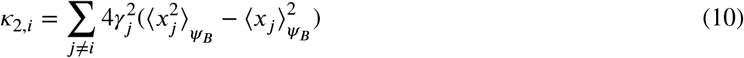

is the *statistical variance* of allele frequency *x*_*i*_ w.r.t. the distribution *ψ*_*B*_ and given by (A3.6). As each summand on the right-hand side (RHS) of (10) is positive, *κ*_2,*i*_ increases linearly with *L* and captures the effect of epistatic interactions, as explained below.

For 2*N sκ*_2,*i*_ ≫ 1, the last factor in (9) reduces to 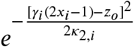, and tends to unity provided *κ*_2,*i*_ is also large (and *z*_*o*_ does not scale linearly with *L*). Thus, when *κ*_2,*i*_ ≫ 1 *and* 2*Nsκ*_2,*i*_ ≫ 1, the effect of epistatic interactions can be ignored and the marginal distribution (9) reduces to *ψ*_*B*_(*x*_*i*_):

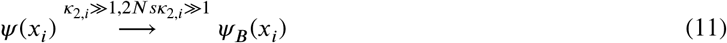

On approximating 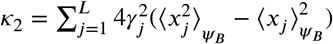, as shown in Appendix A4, we obtain

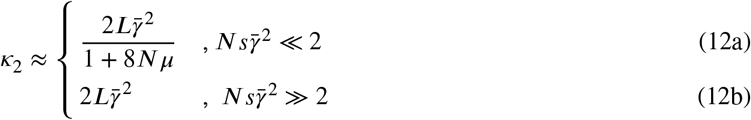

For 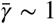, when selection is strong 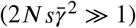, it is sufficient to ensure that 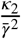 is also large in order to approximate the marginal distribution by *ψ*_*B*_ . Then (12b) shows that the epistatic factor in the marginal distribution can be ignored for large number of loci. However, for weak selection, *κ*_2_ given by (12a) must be much larger than (2*Ns*)^−1^ in order that epistasis can be neglected.

For large *L*, the exponential on the RHS of (9) can be expanded in powers of *L*^−1^; however, as explained in Blinnikov and Moessner (1998), corrections to central limit theorem are required to obtain the correct expression to leading order in *L*^−1^, and is briefly described in Appendix A5.

### 4.2. Transition in marginal distribution for strong mutation

In Fig. 2, we show the marginal distributions obtained in simulations and in Fig. 3 and Fig. 4 by solving the Langevin equation (5) numerically, and compare them with the analytical expression (9). For weak mutation (4 1), as fig. 2 shows, the marginal distribution (9) is U-shaped, as expected. For the parameters in this figure, yields *κ*_2_ ≈ 3.4, 2*Nsκ*_2_ ≈ 344 so that (11) is satisfied, and we observe a good match with *ψ*_*B*_(*x*_*i*_).

**Figure 2.**
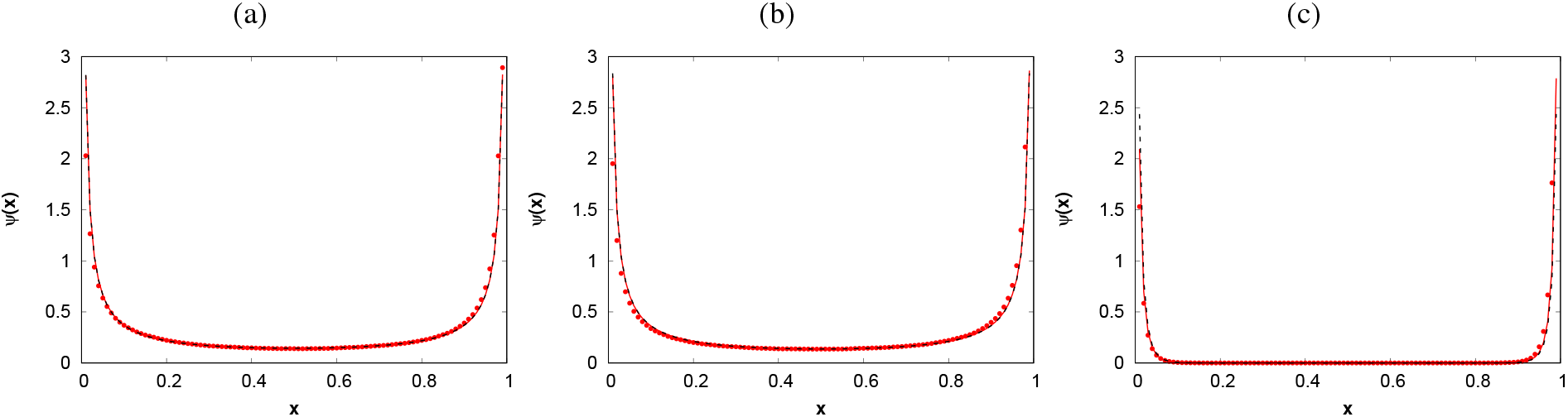
Marginal distribution for weak mutation (4*Nμ* < 1) at a locus with effect size (a) *γ*_*i*_ = 0.01, (b) *γ*_*i*_ = 0.05 and (c) *γ*_*i*_ = 0.7. The parameters are *N* = 1000, *s* = 0.05, *μ* = 0.00002, *L* = 200, 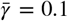, and *z*_*o*_ = 1. The deterministic threshold size 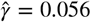 for these parameters whereas stochastically, there is no threshold effect. The points are obtained from MC simulations and the red solid line shows the analytical expression (9) where *κ*_2_ ≈ 4.8 for the effect sizes used in this plot. Here, 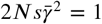 and as both *κ*_2_ and 2*Nsκ*_2_ are large, (11) is satisfied and the marginal distribution (8) shown by black dashed line matches the distribution *ψ*.

**Figure 3.**
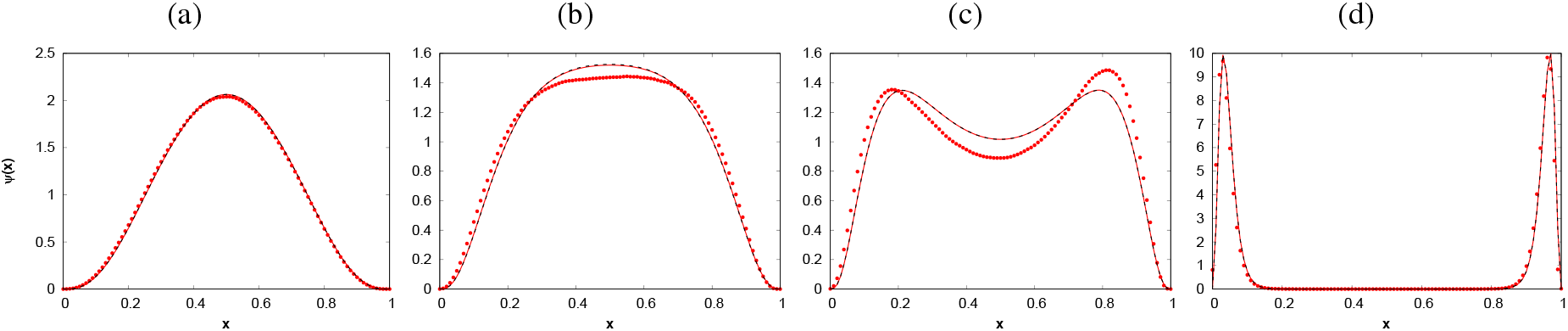
Marginal distribution for strong mutation (4*Nμ* > 1) and strong selection 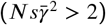 at a locus with effect size (a) smaller (*γ*_*i*_ = 0.1), (b) just below (*γ*_*i*_ = 0.23), (c) just above (*γ*_*i*_ = 0.3) and (d) larger (*γ*_*i*_ = 0.7) than the threshold size 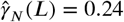. The parameters are *N* = 1000, *s* = 0.1, *μ* = 0.001, *L* = 200, 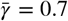, and *z*_*o*_ = 1. For the set of effects used here, there were 137 large-effect loci. The points are obtained by solving (5) numerically, and the red solid line shows the analytical expression (9) where *κ*_2_ ≈ 235 for the effect sizes used in this plot. Here, as both *κ*_2_ and 2*Nsκ*_2_ are large, (11) is satisfied and the marginal distribution (8) shown by black dashed line matches the distribution *ψ*.

**Figure 4.**
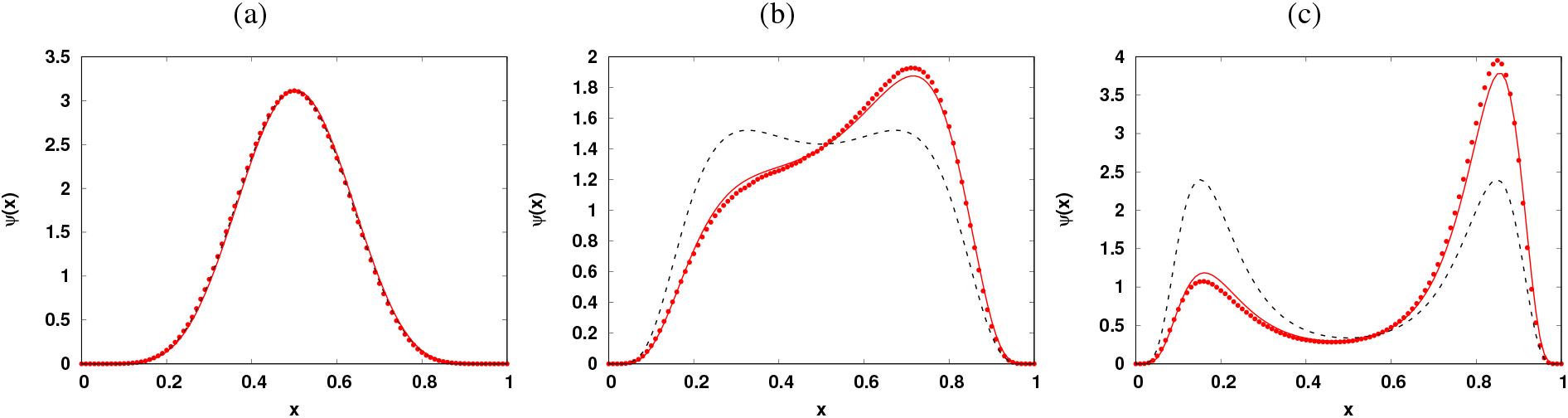
Marginal distribution for strong mutation (4*Nμ* > 1) and weak selection 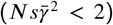 at a locus with effect size (a) smaller (*γ*_*i*_ = 0.05), (b) close to (*γ*_*i*_ = 0.4) and (c) larger (*γ*_*i*_ = 0.5) than the threshold size 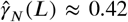. The parameters are *N* = 1000, *s* = 0.1, *μ* = 0.002, *L* = 1000, 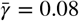, and *z*_*o*_ = 2. For the set of effects used here, there were 7 large-effect loci. The points are obtained by solving (5) numerically, and the red solid line shows the analytical expression (9) where *κ*_2_ ≈ 1.24 for the effect sizes used in this plot. Here, although 2*Nsκ*_2_ ≈ 248 is large, as *κ*_2_ is not very large, (11) is not satisfied and the marginal distribution (8) shown by black dashed line does not match the distribution *ψ* at loci except when the effect size is small.

For strong mutation (4 *N*_*μ*_ > 1), the distribution *ψ*(*x*_*i*_) has the following interesting property: it is unimodal if effect size is below a the threshold effect size 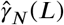 (see Sec. 4.5) and bimodal otherwise. This is displayed in Fig. 3 where the frequency distribution at a locus with sufficiently small effect size has one peak and it is flat for intermediate frequencies at a locus with the threshold size (inflection point), while at a locus with large effect size, the distribution has two maxima which are quite symmetric about frequency one half for parameters in Fig. 3 and asymmetric in Fig. 4. These behavior are also reminiscent of second order phase transitions in physical or chemical systems where the potential energy (given here by − ln *ψ*) changes as an external parameter such as temperature (here, 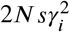) is varied. For the parameters in Fig. 3, as 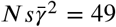, using (12b), we find that *κ*_2_ = 196 which is comparable to value obtained using the effects used in this figure, and as selection is strong and *κ*_2_ is large, we find that *ψ* ≈ *ψ*_*B*_ ; on the other hand, for the parameters in Fig. 4, since 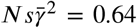, (12a) yields *κ* .75 and as discussed above, the marginal distribution *ψ*is poorly estimated by *ψ*_*B*_ .

In Fig. 3 and Fig. 4, we also note a mismatch between the numerical data and the analytical expression (9) when the locus has an effect size close to or above the threshold size. This is because the population spends a long time near one of the maxima before traversing the valley separating the other maximum (Barton and Rouhani, 1987). Due to such shifts in the equilibria for large-effect loci, the numerical results for the stationary state distribution were obtained by averaging over 103 independent initial conditions (ensemble-averaging) as well as long time periods in the stationary state (time-averaging), as this allowed us to sample the distribution near both the allele frequency peaks efficiently.

### 4.3. Phenotypic mean deviation

Using the joint distribution (6), we find that the distribution of phenotypic mean (7) is given by

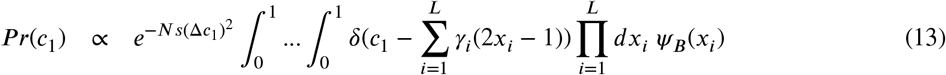

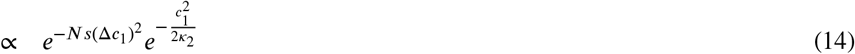

where Δ*c*_1_ = *c*_1_ − *z*_*o*_, *κ*_2_ is given by (A4.2) and the last expression is obtained using the central limit theorem for large *L* (Bulmer, 1972; Lande, 1976). The above distribution gives the average and variance of the phenotypic mean to be

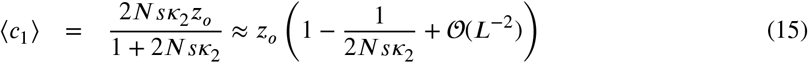

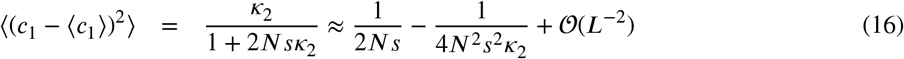

Then the deviation in the trait mean can be written as

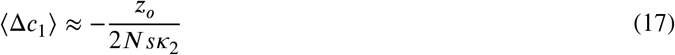

From (15) and (12), we find that if the magnitude of the phenotypic optimum does not increase with the number of loci, the deviation in the mean phenotype vanishes with increasing *L* and the population is perfectly adapted (on an average). Intuitively, as the variance of mean becomes smaller with increasing *L*, the distribution of phenotypic mean gets narrower so that the first factor in (9) (although still random) can be neglected.

### 4.4. Mean genic variance

As discussed in Appendix A1, the genic variance within the population is defined as

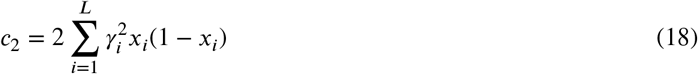

where the mutant allele frequencies are random variables distributed according to (6). For equal effects and *z*_*o*_ = 0, Bulmer (1972) has obtained an expression for the expectation value of the genic variance by arguing as follows: from (6), the expected genic variance can be rewritten as

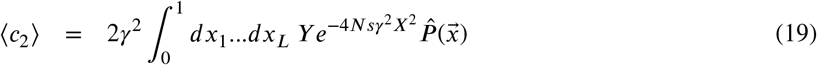

where 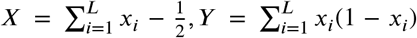 and 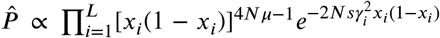 . Then, since *X* is antisymmetric about *x*_*i*_ = 1/2 while *Y* and 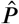 are symmetric, it follows that the expectation value of *XY* with respect to 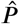 is zero and it is concluded that “Hence *X* and *Y* must also be uncorrelated” (Bulmer, 1972). This reasoning then gives

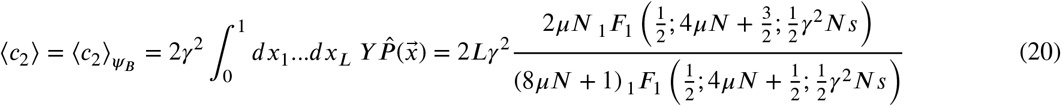

where _1_*F*_1_(a, *b, z*) is the confluent hypergeometric function [Chapter 13, (Olver, Olde Daalhuis, Lozier, Schneider, Boisvert, Clark, Miller, Saunders, Cohl and McClain, 2024)]. However, the exponential factor in the integrand on the RHS of (19) is an even function of *X*, and on expanding the exponential in a power series in *X*, we immediately find that the expectation value of 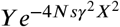 and *Y* w.r.t. 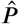 are not equal, contrary to Bulmer’s argument above.

For two loci, the integrals in (19) can be readily done numerically and the expected variance is shown in Fig. 5a and Fig. 5b, and compared with 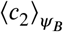. We find that Bulmer’s expression (20) does not match the exact result except for very weak selection (neutral limit) where the fitness epistasis is negligible and for very strong selection where variance in mean deviation given by (16) is negligible so that the loci evolve independently. These results are also compared with those obtained numerically. We find that the mean genic variance ⟨*c*_2_⟩ and mean genetic variance ⟨*C*_2_⟩ (where ⟨*c*_2_⟩ is defined in (A1.3)) obtained in MC simulations are equal since the linkage disequilibrium is negligible for weak to moderate mutation rate with small number of loci where the recombination probability of 0.5 can maintain the linkage equilibrium. The results from MC simulations are also in agreement with the exact results for both weak and strong mutation; however, the corresponding results for genic variance from ⟨*c*_2_⟩ EM method do not match the exact results for weak mutation for the reasons discussed in Sec. 3.

**Figure 5.**
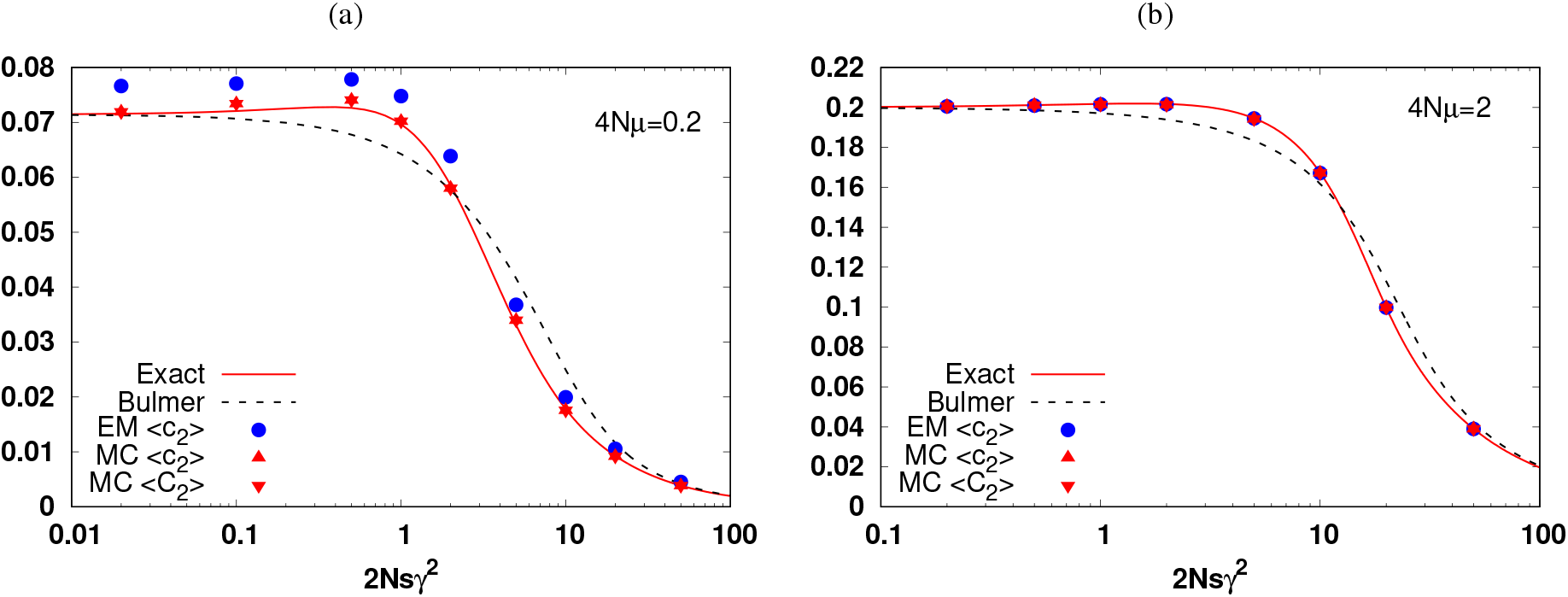
Mean genic variance (18) for *L* = 2 for (a) weak and (b) strong mutation. In all the figures, *N* = 1000, *z*_*o*_ = 0, *γ*_*i*_ = 0.5 (equal effects). In the legend, the line denoted by ‘Exact’ is obtained by numerically integrating the double integral of the joint distribution in the result (19) of diffusion theory, whereas ‘Bulmer’ shows the result (20) obtained by Bulmer, while ‘MC’ and ‘EM’, respectively, represent the data obtained using Monte-Carlo simulations and numerical integration of Langevin equation via Euler-Maruyama method.

In Fig. 6 where genic variance and genetic variance are shown for large *L*, we find that the mean genic variance is close to 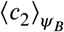. However, the mean genic variance and the mean genetic variance in MC simulations are not equal for strong selection and strong mutation since the interference between genotypes is strong in this regime leading to a negative linkage disequilibrium between loci, and recombination with probability one half can not break all the associations between the loci; this results in a reduction in the genetic variance (Bulmer (1971, 1974)) which depends on the genic variance ⟨*c*_2_⟩ and the fitness variance 1/ (Turelli (1988); Turelli and Barton (1990, 1994)). Figure 6 also shows that the EM method does not work for weak mutations, but it is in closer agreement with Bulmer’s expression than the MC simulation data when mutation is strong. From (9), we find that for large *L*, the corrections to Bulmer’s result are of order *L*−1, and these are discussed in Appendix A6.

**Figure 6.**
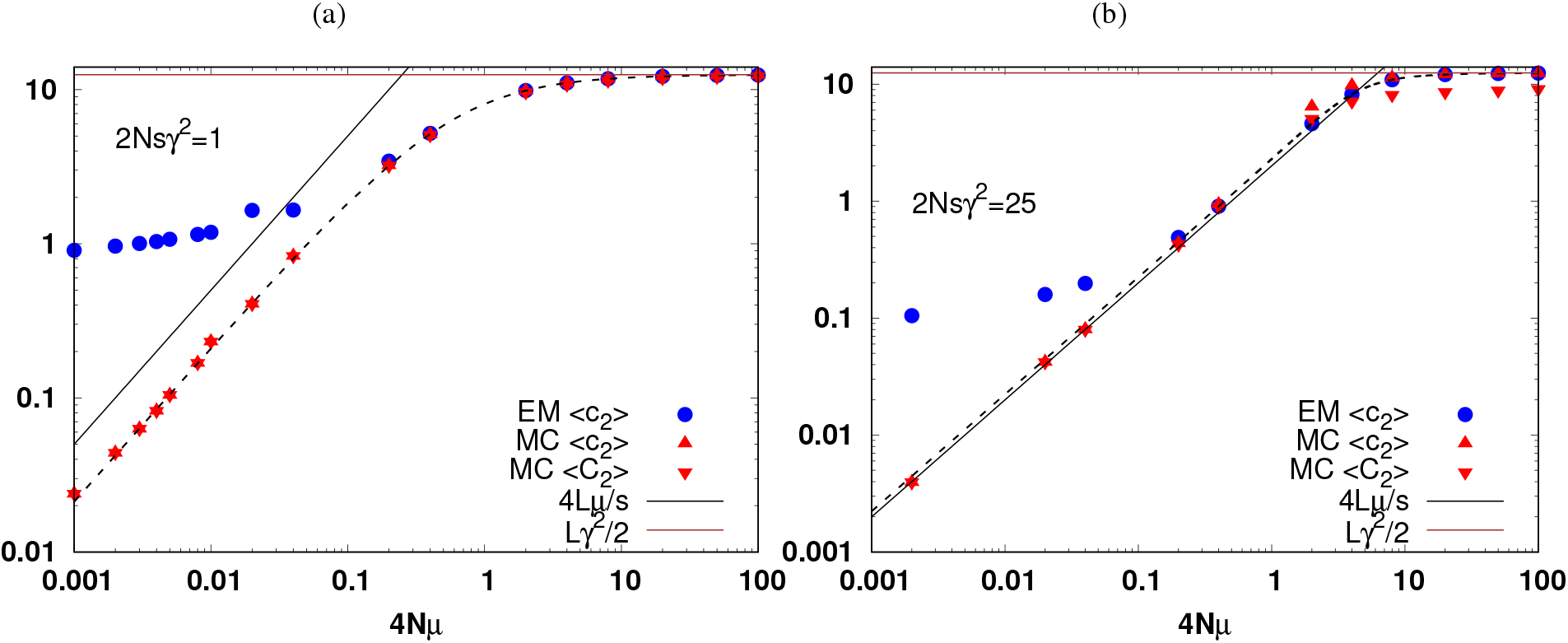
Mean genic variance (18) and mean genetic variance (A1.3) for *L* = 100 for (a) weak and (b) strong selection. In all the figures, *N* = 1000, *z*_*o*_ = 0, *γ*_*i*_ = 0.5 (equal effects). In the legend, the ‘MC’ and ‘EM’, respectively, represent the data obtained using Monte-Carlo simulations and numerical integration of Langevin equation via Euler-Maruyama method. The black dashed line shows the result (20) obtained by Bulmer, while the black solid line shows the genic variance in the rare allele approximation (Barton, 1986), and the brown line shows the genic variance corresponding to all frequencies equal to one half in the strong mutation limit.

### 4.5. Threshold effect size

We finally describe the threshold effect size introduced in Sec. 4.2, and how it depends on *L* and *N*. For *L* → ≈, taking the derivative of the distribution (8) w.r.t. *x*_*i*_, we immediately find that for 4*Nμ* > 1, the allele frequency distribution has one maximum if the effect size at a locus is below the threshold effect size 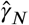 and two maxima otherwise, where 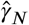 is given by

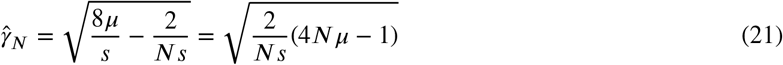

The maxima of the allele frequency distribution (8) occur at

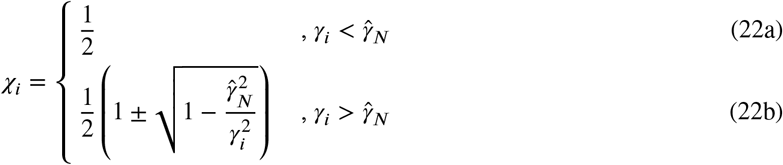

and for 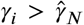, the maxima of the bimodal distribution are separated by a minimum at frequency 1/2. Thus, (21) and (22) show that when selection is weaker than mutation 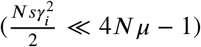 or equivalently, for *small effect size* 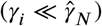, due to a high degree of polymorphism, the allele frequency distribution peaks at one half; however, when selection is stronger than mutation or for *large effect size*, the allele frequency distribution peaks at frequencies away from one half but the population is not monomorphic (unlike the U-shaped distribution in weak mutation regime).

The finite *L* corrections to 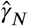 and χ_*i*_ are derived in Appendix A7, and shown in Fig. 7; we find that the modes of the allele frequency distribution at a locus with effect size away from the threshold effect 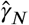 (*L*) are well approximated by the result (22) for infinite loci. But close to the threshold effect, they are substantially different; in particular, for positive (negative) phenotypic optimum, the maximum in the frequency distribution of the small-effect locus increases (decreases) with the effect size and occurs at a frequency which is substantially larger (smaller) than one half.

**Figure 7.**
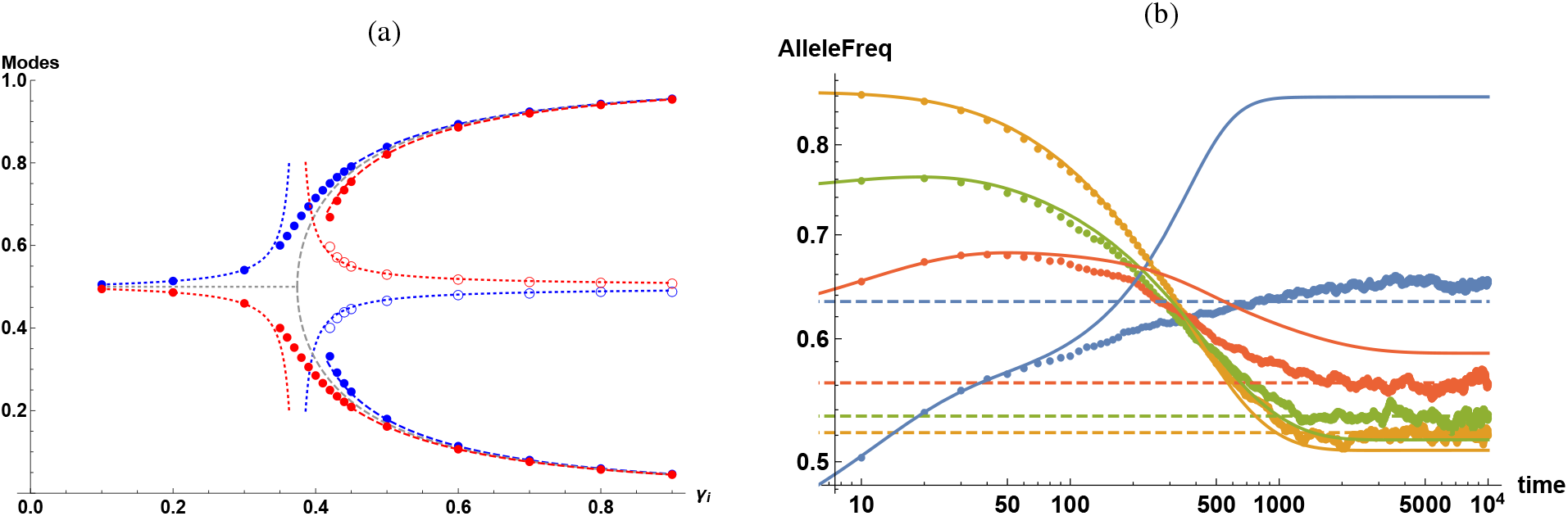
(a) Allele frequency at which the modes in the stationary state marginal allele frequency distribution occur for *z*_*o*_ = 2 (blue) and −2 (red). The points are obtained by numerically solving the cubic equation (A7.1) for the modes, with closed (open) symbols denoting the maximum (minimum). The dotted and dashed lines, respectively, show the approximate expressions in (A7.3a) and (A7.3b) for the mode frequency for finite number of loci while the gray lines show (22) for infinite number of loci. The parameters are the same as in Fig. 4, *viz., N* = 1000, *s* = 0.1, *μ* = 0.002, 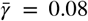, and *L* = 1000. The threshold effect obtained from the cubic equation (A7.2) and the approximation (A7.5) are, respectively, 0.416 and 0.401, while (21) yields 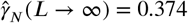. (b) Dynamics of deterministic allele frequency (solid lines) obtained by numerically solving (23) and average allele frequency (points) obtained by numerically solving (5) for 5000 independent stochastic runs for effect size 0.24 (yellow), 0.36 (green), 0.52 (red), 0.8 (blue), keeping the initial frequencies to be the same in both deterministic and stochastic model. In either model, the population is initially equilibrated to an optimum at *z*_*o*_ = 0 and then the optimum is suddenly shifted to *z*_*o*_ = 1 in response to which the allele frequencies evolve in time. The other parameters are *L* = 200, *s* = 0.05, *μ* = 0.002, *N* = 200. The threshold effect 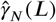 is 0.359 and 0.493 when the optimum is at zero and one, respectively. The dashed lines show the average allele frequency *x* in the stationary state for phenotypic optimum equal to one, and are computed using the marginal distribution (9).

It is also useful to compare these results for an infinitely large population where the deterministic allele frequency *p*_*i*_ obeys (de Vladar and Barton, 2014; Jain and Stephan, 2017b)

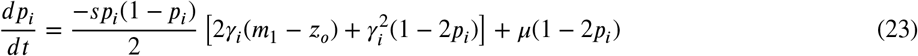

with the deterministic trait mean, 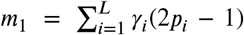. Assuming that the trait mean deviation is zero in the stationary state, it has been shown that the deterministic allele frequency is in stable equilibrium below a threshold effect 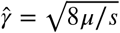 and is bistable above it (de Vladar and Barton, 2014), and given by

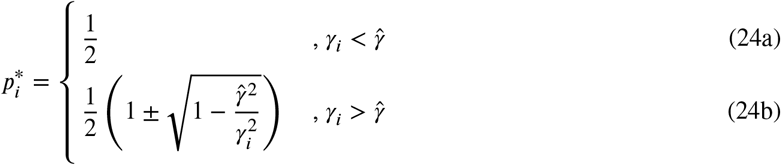

## 5. Discussion

In this article, we considered a model that describes a large but finite population under stabilizing selection and symmetric mutation in the equilibrium state. The dynamics of adaptation following a sudden change in the phenotypic optimum of such a model have been studied in deterministic (de Vladar and Barton, 2014; Jain and Stephan, 2015, 2017b) and stochastic setting (Thornton, 2019; Hayward and Sella, 2022; Höllinger, Wölfl and Hermisson, 2023).

Here we asked: in what parameter regimes the fitness epistasis can be neglected when the population is in equilibrium? Our key result for the allele frequency distribution is summarized in (11). We find that for strong selection, it is sufficient to have large number of loci in order to describe the distribution without taking epistasis into account, but for weak to moderate selection, there are conditions on mutation and selection parameters given by (12) under which epistasis can be neglected.

To connect these results at the genetic level to the phenotypic quantities, we also studied the phenotypic deviation and mean genic variance. We find that Bulmer’s argument (Bulmer, 1972) for mean genic variance is incorrect and therefore, their expression given by (20) is not exact, but it is a good approximation when the number of loci are large. We stress that the disagreement with Bulmer’s expression is not because of linkage disequilibrium [Bulmer effect; (Bulmer, 1971)] as the diffusion theory formulated here is in linkage equilibrium rather the point is that (20) neglects the epistatic effects. The discussion in Bulmer (1972) does not invoke the number of loci, and we are not aware of other work where this error has been pointed out. Thus, for small number of loci, as in oligogenic traits (Bell, 2009; Boyle, Li and Pritchard, 2017), the expression (29) of Bulmer (1972) does not hold but it is a good approximation for large *L* for any selection strength.

Taken together, our results show that for weak to moderate selection, although genic variance matches well with Bulmer’s expression, epistasis strongly affects the allele frequency distribution at loci with large effect size. Thus, while epistasis effects may not be evident at the phenotypic level, they can strongly influence the distribution at the genetic level. In a recent study by Courau, Lambert and Schertzer (2026), the model here has been generalized to include heterogeneous mutation rates and asymmetric mutations, and the authors arrive at the same general conclusions as in this article but via a different route.

We close the article by discussing the key differences in the stationary state of infinite (de Vladar and Barton, 2014; Jain and Stephan, 2017b) and finite populations: first, it should be noted that, in general, there is no one-to-one correspondence between the multistability in a deterministic model and multimodality in the corresponding stochastic model. This point, for example, has been made in the context of biochemical reaction systems where some monostable (bistable) deterministic systems have been observed to have a bimodal (unimodal) distribution when stochastic fluctuations are allowed (Hahl and Kremling, 2016). Here, as 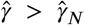, when 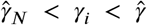, the deterministic allele frequencies are monostable but the allele frequency distribution is bimodal; however, outside this parameter range, the bistability (monostability) in the deterministic model corresponds to bimodality (unimodality) in the stochastic model.

Second, the stationary state solution of the Fokker-Planck equation (A2.1) is unique [refer to Chapter 5, (van Kampen, 1997)]; that is, it is independent of the initial allele frequencies for both small- and large-effect loci. Furthermore, as already mentioned, in a finite population, the allele frequency of a large-effect locus spends a long time (presumably, exponentially long in population size) near one of the maxima before shifting to the other maximum. In contrast, in an infinite population, the stationary state allele frequency of a large-effect locus depends on the initial condition and does not shift between the two solutions given in (24b). These points are further illustrated in Fig. 7b where the stationary state frequencies in the deterministic and stochastic model, starting from the same initial condition, are found to be quite close for small effect loci but not for the large-effect ones. For the latter case, to obtain a match between the *N* → ∞ limit of the stochastic model and the deterministic model, one also need to average over the initial conditions in the deterministic model.

### A1. Linkage equilibrium

Let *f*_*σ,σ*_∗ denote the frequency of a diploid individual with maternal and paternal sequences, *σ* and *σ*^∗^, respectively. If the phenotypic effect size at locus *i* is *γ*_*i*_/2, for equivalent sexes and assuming additive phenotype-genotype map, the trait value of the individual can be written as

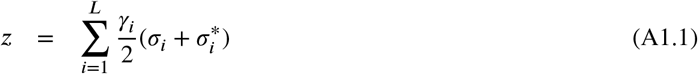

with *σ*_*i*_ = 1(−1) for mutant (wildtype) allele. Under random mating, the parent frequencies factorize in Hardy-Weinberg equilibrium (HWE). Then the mean phenotype of the population (denoted by bar) obtained by averaging over the phenotypes of all individuals is given by

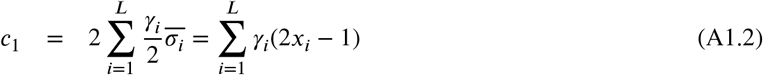

where 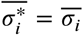 for equivalent sexes, and *x*_*i*_ is the frequency of the mutant allele at locus *i*. Similarly, one obtains the within-population (additive) genetic variance as (Bulmer, 1971)

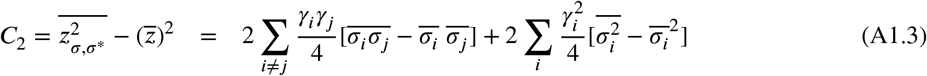

Here, we assume that the population is in linkage equilibrium, 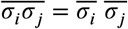,and obtain the (additive) genic variance, *c*_2_ given by (18).

## A2. Diffusion theory

Let 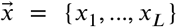 denote the mutant allele frequency vector and 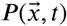 is the joint distribution of these frequencies. The forward Fokker-Planck equation (FPE) is then given by

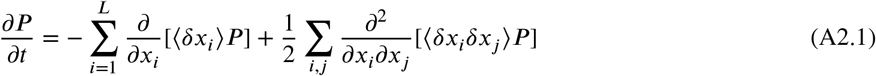

where, the mean, variance and covariance in the change in the allele frequencies due to stabilizing selection, mutation and random genetic drift are described below.

First consider the effect of selection: Since 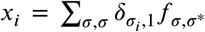, the (deterministic) change in the mutant allele frequency after selection is given by (Kimura, 1964; Bulmer, 1972)

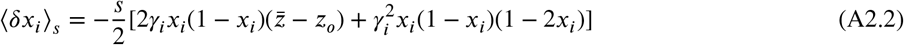

on using that the population is in HWE before selection. For symmetric mutation rates, the deterministic change in *x*_*i*_ due to mutation is given by

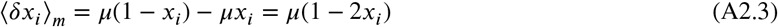

Thus the change ⟨*δx*_*i*_⟩ due to selection and mutation is obtained by adding their respective contributions, and given by

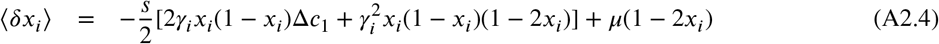

Note that ⟨*δx*_*i*_⟩ depends on the frequency of other loci also through 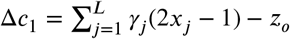 . Finally, the change due to multinomial sampling in the Wright-Fisher process is given by

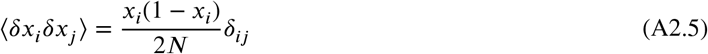

where the covariance vanishes as the population is in linkage equilibrium.

In the steady state [refer to Chapter 6, (Risken, 1996); (Kimura, 1964)], the LHS of (A2.1) is zero, and therefore, the divergence of the total probability current vanishes: 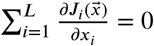, where

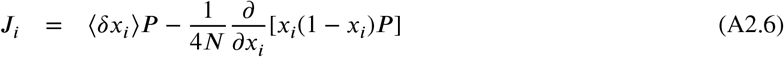

In general, it is not necessary that each *J*_*i*_ be same for all loci, and moreover, equal to zero. But if we do assume that *J*_*i*_ = 0 for all *i*, we obtain

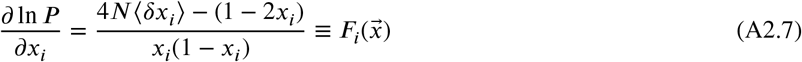

Then, for Φ = ln *P*, the total derivative 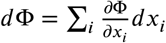 exists iff 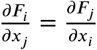; using (A2.4), we verify that this condition is indeed satisfied, and therefore, we can write

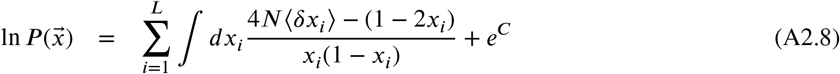

We thus obtain

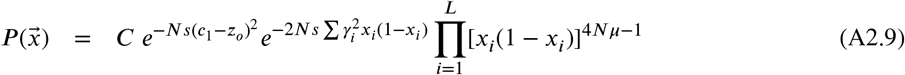

where, the constant *C* is determined using the normalization condition, 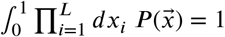.

## A3. Marginal distribution

The marginal distribution *ψ*(*x*_*i*_) of the allele frequency in the stationary state can be found by integrating the joint distribution 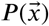 given by (6) over the frequencies of all but the *i*th locus; this gives

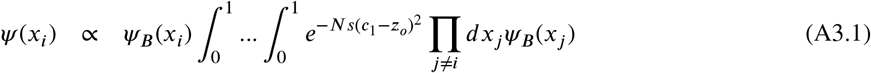

and 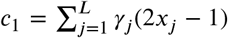. To evaluate the multiple integrals in (A3.1), we take 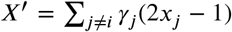 and rewrite its RHS as

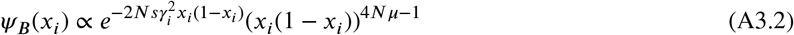

where Γ′ = ∑_*j*≠*i*_*γ*_*j*_, and *X*′ is a sum of independent but non-identically distributed random variables chosen from *ψ*_*B*_(*x*_*j*_). The inner integrals on the RHS of the above equation can be calculated by appealing to the central limit theorem for large *L*, and we get

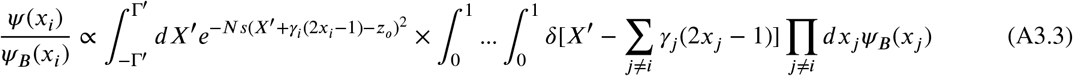

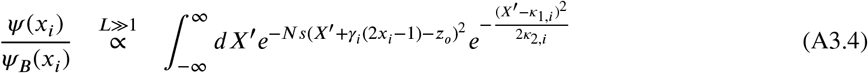

where *κ*_1,*i*_ and *κ*_2,*i*_ are, respectively, the mean and variance of the sum *X*′= ∑ _*j*≠*i*_ *γ*_*j*_ (2*x*_*j*_ − 1) when averaged over the (normalized) distribution ∏_*j*≠*i*_ *ψ*_*B*_(*x*_*j*_).

Since *ψ*_*B*_(*x*_*i*_) is symmetric about *x*_*i*_ = 1/2, it follows that the mean *κ*_1,*i*_ = 0. In the marginal distribution *ψ*(*x*_*i*_), the effect of other *L* − 1 loci due to epistatic interactions in the phenotypic fitness function appears through the variance *κ*_2,*i*_. Since 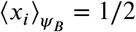 for all loci, the variance 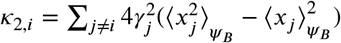 which shows that *κ*_2,*i*_ is simply a weighted sum of the variance of the distribution *ψ*_*B*_ (*x*_*i*_), and is given by

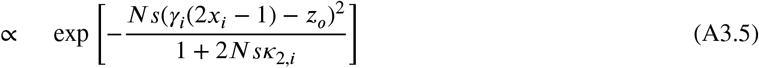

where _1_*F*_1_(*a, b, z*) is the confluent hypergeometric function.

## A4. Statistical variance in the absence of epistasis

Here, we study the properties of 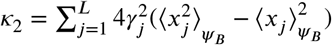 . Note that *κ*_2_ is the standard statistical variance while the mean genic variance 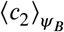 is related to the expectation of *x*(1 − *x*), although both are calculated w.r.t. *ψ*_*B*_. We have

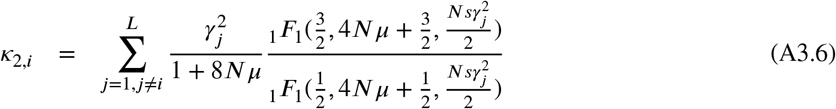

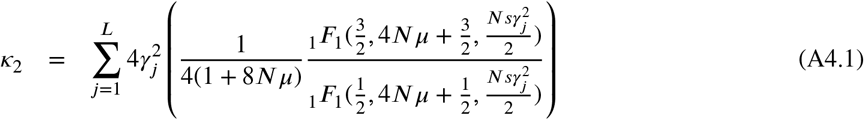

where *p*(*γ*) is the distribution of effects. For exponentially-distributed effect size with mean 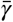, we have

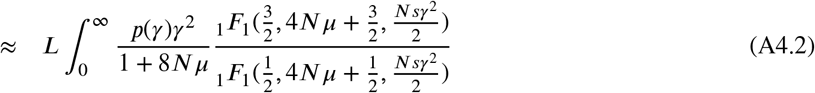

As the above integral does not appear to be exactly solvable, we estimate it by noting that the ratio of the confluent hypergeometric function in the above integrand is a monotonically increasing function of its argument, and for *fixed U* = 4N*μ*, this ratio may be approximated using (13.7.1) and (13.11.1) of Olver et al. (2024) as

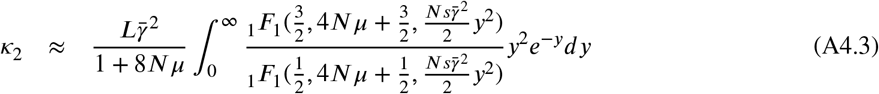

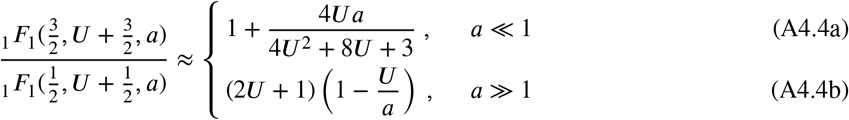

Thus, for a given 4*Nμ*, using (A4.4) in (A4.3), we obtain

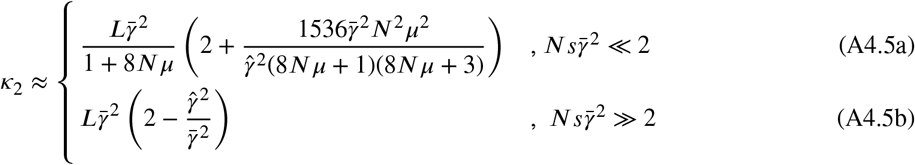

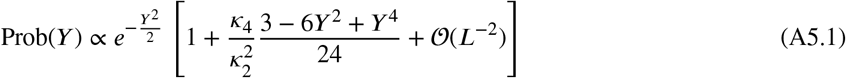

where,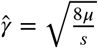.

When selection is weak 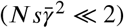 and mutation is strong (4*Nμ* ≫ 1), (A4.5a) shows that *κ*_2_ approaches zero as 1/*N*; in contrast, when selection is stronger than mutation, due to (A4.5b), the variance 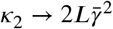 so that it remains nonzero in the deterministic limit. These results can be understood as follows: as the width of *ψ*_*B*_ (*x*_*i*_) about a maximum is expected to decrease with *N*, for large *N*, one may approximate *ψ*_*B*_(*x*_*i*_) by a Dirac delta function centred at 1/2 for small effect locus, and an average of two Dirac delta functions located at 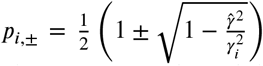 for large-effect locus (de Vladar and Barton, 2014). Thus, for small-effect locus, as the distribution is unimodal and sharply-peaked in large populations, the variance vanishes in the deterministic limit. But for large-effect locus, as a consequence of the bimodality, the distribution remains broad resulting in a nonzero variance.

## A5. Corrections to central limit theorem

Consider the distribution of the random variable, 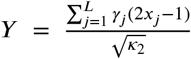 where *x*_*i*_’s are independently distributed according to the (normalized) distribution, 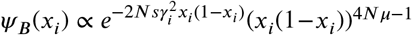 and 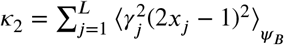 where we have used that the mean 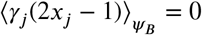 . Then, from (43) of Blinnikov and Moessner (1998), we have

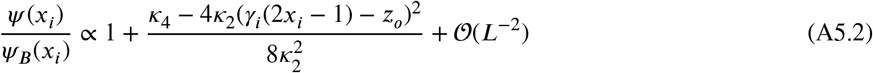

where, 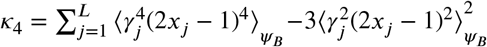 is the fourth cumulant of the random variable *γ*_*j*_(2*x*_*j*_ −1) w.r.t. the (normalized) distribution *ψ*_*B*_ (*x*_*j*_ ). Using the above result in (A3.3) for 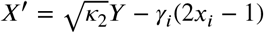 and performing the integral over *X*′, we arrive at

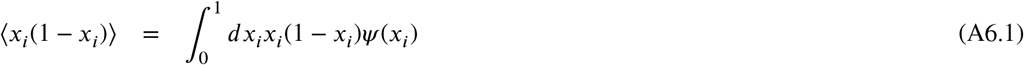

## A6. Stationary state genic variance

Consider the genic variance (18) when it is averaged over the stationary state distribution (A5.2). For *z*_*o*_ ≪ *L*, we can write

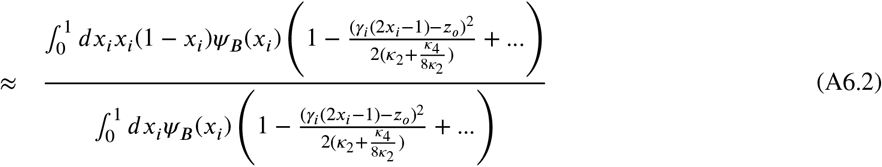

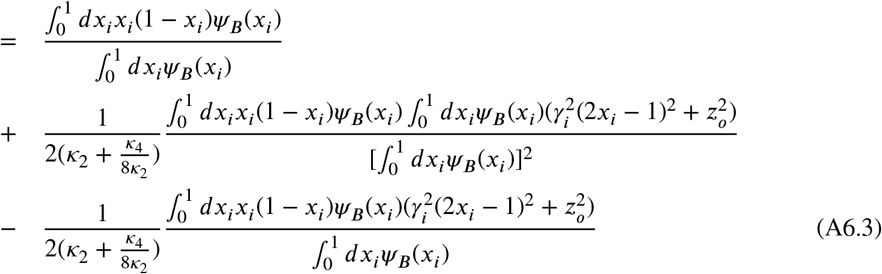

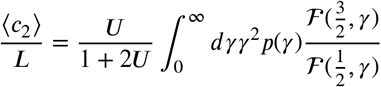

On summing over all the loci, the last equation gives the mean genic variance to be

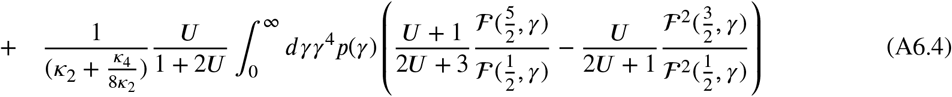

where, for brevity, we have defined 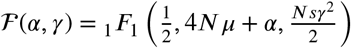 and *U* = 4N*μ*.

The first term on the RHS of (A6.4) is independent of *z* _*o*_ and is equal to ⟨*c*_2_⟩_*ψB*_, and the second term that gives the *L*-dependent correction is of order *L*^−1^ and is also seen to be independent of the phenotypic optimum. Thus, when *z*_*o*_ ≪ L, the mean genic variance (to 𝒪 (*L*^−1^)) does not depend on the location of the phenotypic optimum.

## A7. Threshold effect size and mode frequency

On setting the derivative 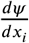 of the marginal distribution (9) equal to zero, we arrive at the following cubic equation for the modes χ_*i*_(*L*):

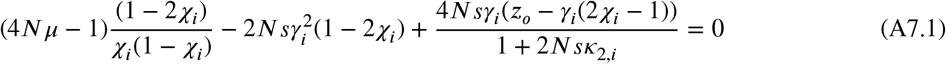

The above equation has two complex roots and one real root below the threshold effect 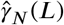 and three real roots above it. This change in the behavior of χ_*i*_(*L*) occurs when the discriminant of the above cubic polynomial [(1.11.12), (Olver et al., 2024)] is equal to zero:

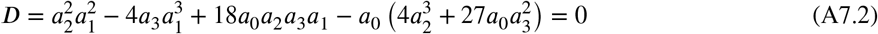

where *a*_*i*_, *i* = 0, 1, 2, 3 are the coefficients of *x*^*i*^ in (A7.1). The resulting equation is a 3rd order equation in 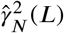 and can be solved numerically to obtain the threshold effect for finite *N* and *L*.

One can, however, obtain an analytical expression for 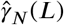 when the number of loci are very large. Since *κ*_2_ grows linearly with *L*, for infinite number of loci, the last term on the LHS of (A7.1) vanishes yielding (21) and (22) for the threshold effect 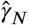 and mode allele frequency χ_*i*_, respectively, which are independent of *z*_*o*_. For large *L*, using (A3.5), we find that the steady state distribution has maximum at

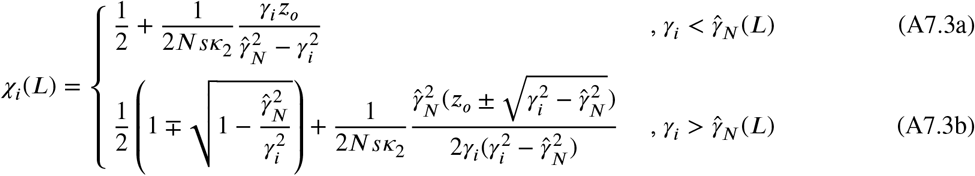

where 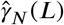 denotes the threshold effect for finite *L*. For 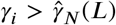, the minimum in the allele frequency is given by the expression on the RHS of (A7.3a).

As shown in Fig. 7, a threshold effect exists below which (A7.1) has only one real root and corresponds to the maximum in the unimodal distribution. But, above the threshold effect, two additional real roots of (A7.1) appear which give the allele frequency at which the minimum and the second maximum of the bimodal distribution occur. Thus, at the threshold effect, for positive (negative) *z*_*o*_, the minimum and the low-frequency (high-frequency) maximum of the bimodal distribution coincide. On matching (A7.3a) and the relevant solution in (A7.3b), we get

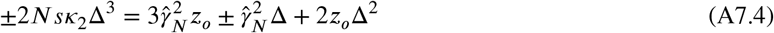

where 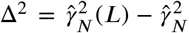 decreases with increasing *L*. The above cubic equation for Δ is exactly solvable, and has two complex ro^*N*^ots and o^*N*^ne real root. Here, we estimate the real root by noting that the first term on the RHS which is independent of *L* can be balanced if *κ*_2_Δ^3^ is also independent of *L* thus yielding

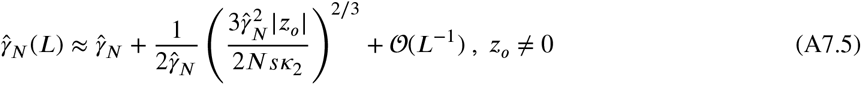

which shows that the deviation 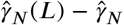 decays rather slowly with *L*. The above expression also shows that the threshold effect always increases with the absolute value of the phenotypic optimum. But it does not change if the phenotypic optimum shifts between *z*_*o*_ and −*z*_*o*_. The threshold effect is also larger when selection is weaker 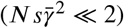 or the quantitative trait is controlled mostly by small-effect loci. For phenotypic optimum at zero, (A7.4) gives

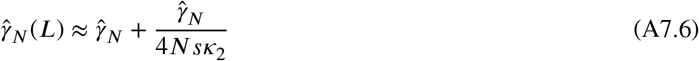

so that the deviation 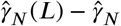 is of order *L*^−1^.

For small *z*_*o*_ and large *L*, (A7.5) shows that the threshold effect does not differ much from the infinite-loci result (21) when selection is strong 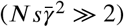. Furthermore, (A7.5) predicts that the threshold effect for large but finite number of loci is always larger than 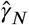, and increases with the magnitude of the phenotypic optimum. Thus, a locus classified as a large-effect locus for a phenotypic optimum at zero can become a small-effect locus if the phenotypic optimum is large enough. This is because for large, positive (negative) phenotypic optimum, population will be well-adapted if the + (−) allele’s frequency at most loci is close to fixation (that is, the distribution is unimodal and heavily skewed towards high frequency).

## CRediT authorship contribution statement

**Archana Devi:** Conceptualization, Formal analysis, Investigation, Methodology, Software, Visualization, Writing - review and editing. **Kavita Jain:** Conceptualization, Formal analysis, Investigation, Methodology, Supervision, Writing - original draft, Writing - review and editing.

